# Bridging the translational gap: Implementation of multimodal small animal imaging strategies for tumor burden assessment in a co-clinical trial

**DOI:** 10.1101/462861

**Authors:** S.J. Blocker, Y.M. Mowery, M. D. Holbrook, Y. Qi, D.G. Kirsch, G.A. Johnson, C.T. Badea

**Affiliations:** Center for In Vivo Microscopy, Duke University Medical Center, Durham, NC 27710; Department of Radiation Oncology, Duke University Medical Center, Durham, NC, 27710; Department of Pharmacology & Cancer Biology, Duke University Medical Center, Durham, NC, 27710

## Abstract

In designing co-clinical cancer studies, preclinical imaging brings unique challenges that emphasize the gap between man and mouse. Our group is developing quantitative imaging methods for the preclinical arm of a co-clinical trial studying immunotherapy and radiotherapy in a soft tissue sarcoma model. In line with treatment for patients enrolled in the clinical trial SU2C-SARC032, primary mouse sarcomas are imaged with multi-contrast micro-MRI (T1 weighted, T2 weighted, and T1 with contrast) before and after immune checkpoint inhibition and pre-operative radiation therapy. Similar to the patients, after surgery the mice will be screened for lung metastases with micro-CT using respiratory gating. A systems evaluation was undertaken to establish a quantitative baseline for both the MR and micro-CT systems against which others systems might be compared. We have constructed imaging protocols which provide clinically-relevant resolution and contrast in a genetically engineered mouse model of sarcoma. We have employed tools in 3D Slicer for semi-automated segmentation of both MR and micro-CT images to measure tumor volumes efficiently and reliably in a large number of animals. Assessment of tumor burden in the resulting images was precise, repeatable, and reproducible. Furthermore, we have implemented a publicly accessible platform for sharing imaging data collected during the study, as well as protocols, supporting information, and data analyses. In doing so, we aim to improve the clinical relevance of small animal imaging and begin establishing standards for preclinical imaging of tumors from the perspective of a co-clinical trial.

## Introduction

The successful design and implementation of a cancer clinical trial faces many challenges, not the least of which is the translational relevance of related preclinical findings (1). While invaluable to cancer research, animal studies are typically completed before patient trials, effectively separating them from clinical observations and preventing bidirectional flow of information between the preclinical and clinical studies (2, 3). Further, preclinical protocols often fall short of mimicking clinical criteria, including schedule constraints and procedure time. To address such shortcomings, the implementation of co-clinical trials is of growing interest in cancer research (4, 5). Referring to the concerted execution of analogous animal and patient studies, co-clinical trials provide a setting in which clinical observations can influence the methodology used in animal experiments (6-8). In turn, novel findings from the preclinical arm can inform the patient study. In this way, co-clinical trials represent a strategy for dynamic integration of animal and patient studies to streamline cancer research efforts. However, the opportunity to lessen the translational divide offered by the co-clinical approach can be stifled by the complexities of designing a robust animal study. This is particularly true when considering the incorporation of small animal imaging into co-clinical studies of cancer.

Small animal imaging for co-clinical cancer trials represents an opportunity to simulate better clinical practice in animals (as imaging is often standard of care and essential for assessing disease response in clinical oncology), as well as to expand the information gathered during study. Technologies such as magnetic resonance (MR) can deliver high-resolution images that describe tumor morphology and composition, as well as how tumors change over time or with treatment (9-11). Computed tomography (CT) provides an efficient, non-invasive method to detect new or metastatic lesions prior to development of symptoms (12, 13). The ability to non-invasively image animals at multiple time points greatly enhances the interpretation of tumor progression and/or therapeutic response, providing an avenue for comparison with longitudinal data from patients on study.

As with all translational research, care must be taken to reduce and acknowledge the differences between animal models and human cancer. Investigators must weigh factors such as tumor model selection, disease progression and intervention timelines, the immune condition of the animal, and the metabolic consequences of therapeutic interventions (2, 14, 15). Basing decisions on clinical standards and observations can promote the translation of findings between the arms of a co-clinical trial. However, the challenges that face translational imaging include both the biological and physical differences of the subjects and scanners, respectively.

While protocols for preclinical imaging are often designed with the intention of discovery, they routinely sacrifice efficiency and throughput for innovation (16). These scan programs are commonly project- and machine-specific, limiting their dissemination and broad employment. For example, a brief review of the literature discussing MR imaging in mice to measure tumor volume reveals a great deal of diversity among sequences, and acquisition parameters (Table I). By contrast, clinical scan protocols for tumor imaging are more standardized, achievable on a variety of systems, and typically limit scan durations for patient comfort and to accommodate scanner schedules.

**Table I.**
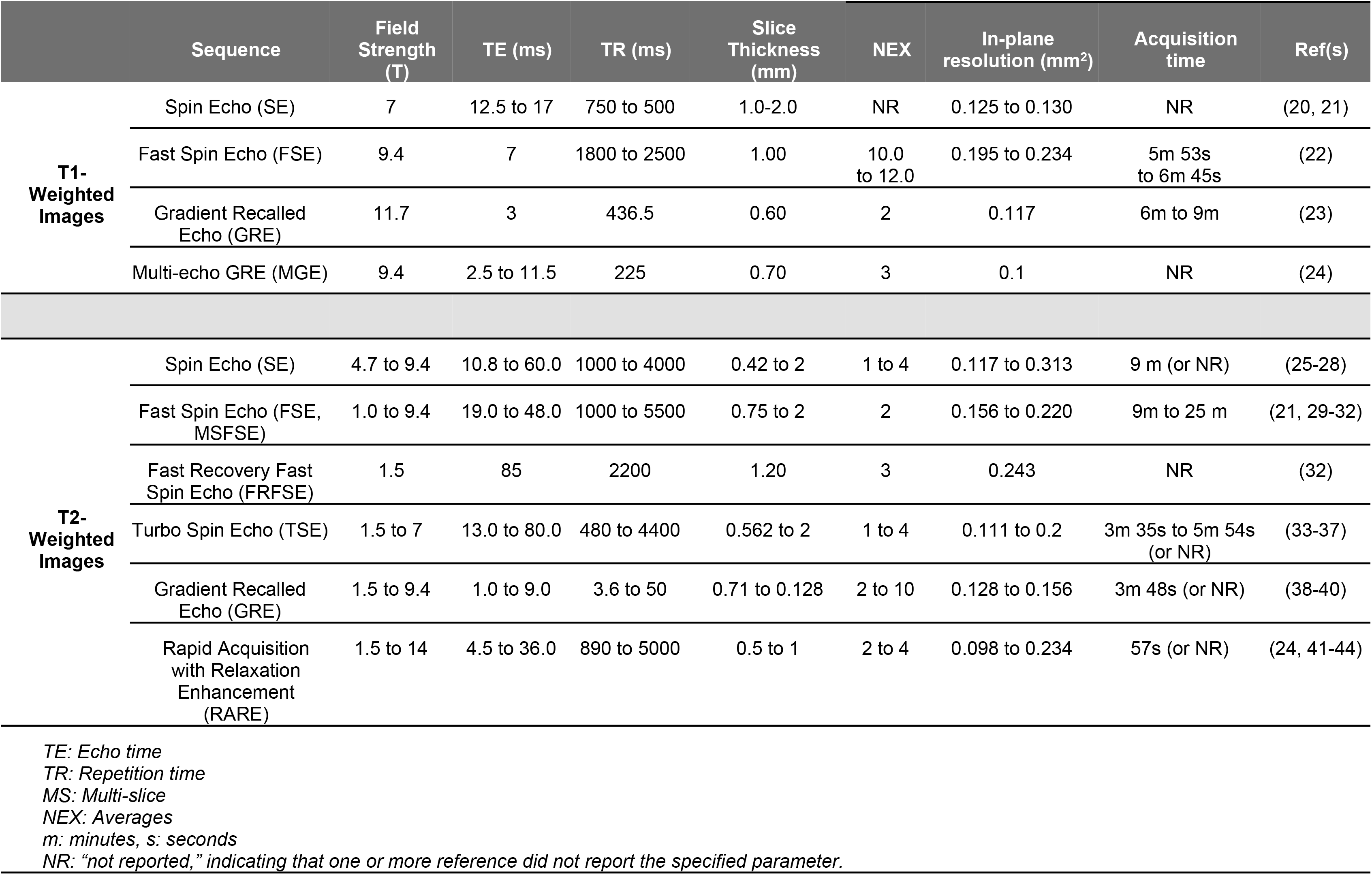
Samples found in the literature on preclinical MR sequences for solid tumor volume measurements in mice

Preclinical scanners must image at much higher resolution than clinical scanners, requiring different hardware solutions. Preclinical MR scanners employ smaller radio frequency coils and often operate at higher field strengths than clinical MR machines (3.0-11.0 T versus 1.5-3.0 T, respectively). The differences in image signal, resolution, artifacts, etc. must all be considered when designing animal imaging protocols to serve a co-clinical trial (17, 18). Thus, the disparity between preclinical and clinical cancer imaging makes correlation of results challenging, potentially reducing the impact and utility of preclinical imaging.

With these challenges in mind we sought to identify clinically-applicable scan protocols to measure tumor burden in a genetically engineered mouse model (GEMM) of soft tissue sarcoma. Specifically, the co-clinical trial requires reproducible detection of changes in primary tumor volume with MR after therapeutic intervention, with follow up detection and measurement of lung metastases with CT. Our goal was to produce images that were comparable to scans performed in an ongoing clinical trial of sarcoma (NCT03092323), while establishing protocols for acquisition, data processing, and image analysis that are reproducible and broadly relevant. These protocols are currently being employed in an ongoing co-clinical trial to assess the effects of neoadjuvant and adjuvant programmed cell death protein 1 (PD-1) inhibition in sarcomas treated with neoadjuvant radiation therapy and surgical resection (19) (see Fig 1). Finally, we have established a resource for public dissemination of preclinical imaging data, protocols, and results. In doing so, we have developed a robust blueprint for incorporating clinically-driven mouse imaging into a co-clinical trial, and a pipeline to promote rigorous reporting and sharing of preclinical imaging practices.

**Fig 1.**
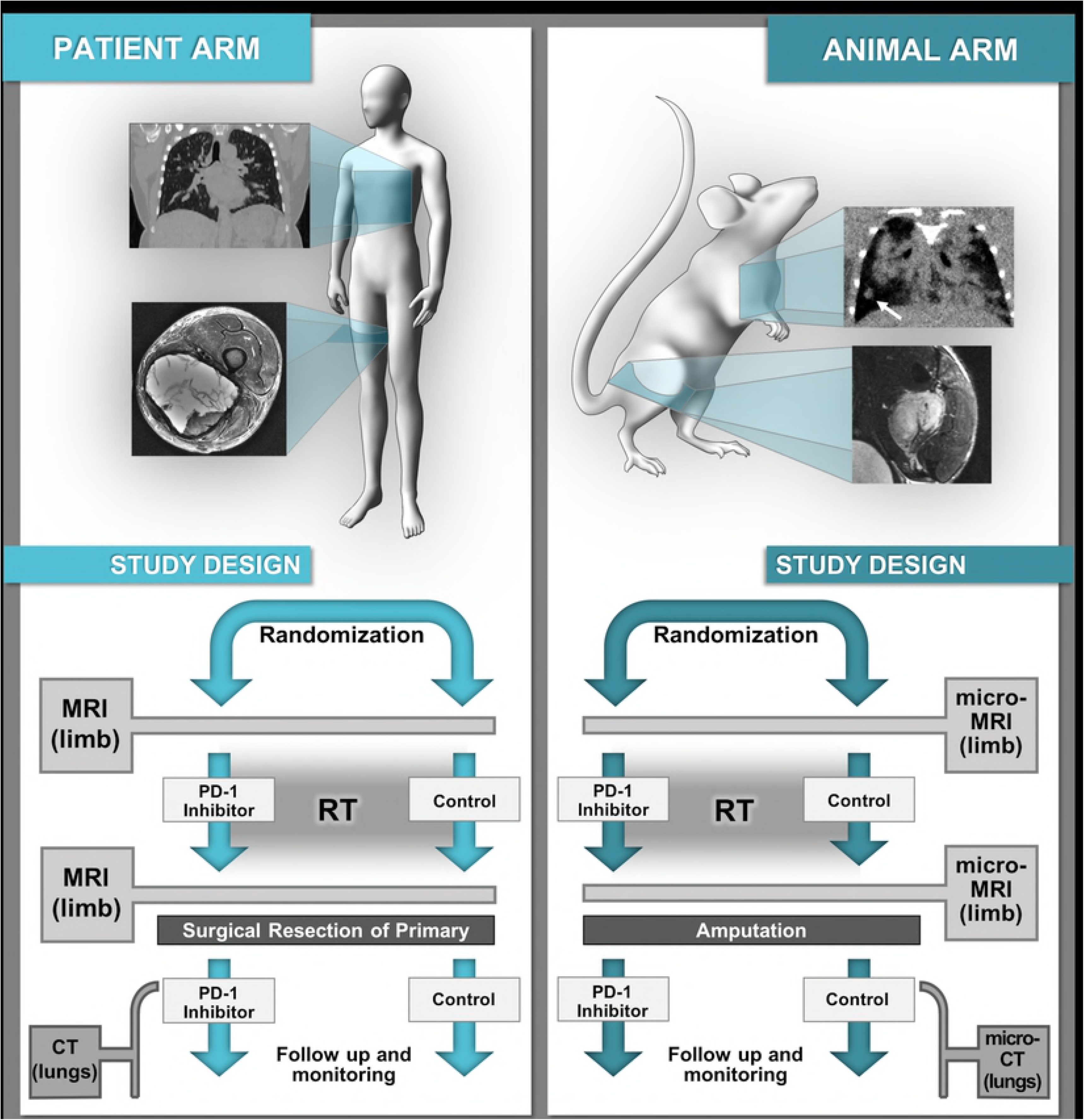
Schematic representation of a co-clinical trial which utilizes translational imaging. Flow charts describing the clinical (A) and preclinical (B) sections of a co-clinical trial of RT with or without the addition of PD-1 inhibition in soft tissue sarcoma. Treatment dosing and imaging procedures in the preclinical arm have been designed to mimic the clinical trial as closely as possible.

## Materials and methods

### GEMM of inducible sarcoma

To mimic the clinical presentation and progression of human sarcomas with gradual tumor development and metastasis in the setting of an intact immune system, a carcinogen-induced GEMM of sarcoma (Lee CL, Mowery YM, Daniel AR, et all, submitted) was chosen for the animal studies of the co-clinical trial. Primary sarcomas (*p53*/MCA model) are generated by intramuscular delivery into the gastrocnemius of adenovirus expressing Cre recombinase (Adeno-Cre; Gene Transfer Vector Core, University of Iowa) into *p53fl/fl* mice followed by intramuscular injection of 0.3 mg 3-methylcholanthrene (MCA; Sigma-Aldrich, Saint Louis, MO) at the same site. Tumors develop approximately 8-12 weeks after induction. Imaging studies were initiated when tumors were palpable (>100 mm^3^), as well as at various stages of disease progression.

### Small animal MR imaging at 7T

All MR experiments were performed on a 7.0 T Bruker Biospec small animal MRI scanner (Bruker Inc., Billerica, MA). In preliminary experiments three commercial Bruker radiofrequency (RF) coils were explored: i) a 72 mm diameter actively decoupled linear volume coil for transmission and reception (Bruker #: T10720V3); ii) a 35 mm diameter quadrature volume coil for transmission and reception (Bruker #: T9988V3); iii) a mouse brain receive-only four coil array (Bruker #: T11765V3), combined with the 72 mm volume coil for transmission. Scan acquisition was performed using ParaVision^®^ 6.0.1 platform (Bruker Inc.), and the reconstructed images were stored in DICOM format.

#### MR System Baseline Performance

A systems evaluation was undertaken to establish a quantitative baseline for our MR system against which others systems might be compared. First, a cylindrical phantom (62 mm diameter, 120 mm length) containing a 3D-printed grid of 1.0 mm squares was scanned with the 72 mm volume coil to measure linearity and geometric distortions across the active volume of the scanner. A second scan was performed of an 18mm x 18 mm cylindrical water bottle filled with 10 M CuSO_4_ to measure B0 homogeneity across the field of view. These data were stored for periodic Quality Assurance (QA) checks during the protocol.

To provide guidance on design of a T1/T2 phantom, a mouse bearing a primary soft tissue sarcoma of the hind limb was scanned in the 72 mm volume coil to obtain T1 and T2 maps across the tumor. The volume coil was used to provide uniform B1 excitation to facilitate accurate T1/T2 mapping. The lower sensitivity of this coil was offset by increasing the number of averages and slice thickness (NEX = 2; Slice Thickness = 1.0 mm). T1 maps were acquired using Multi-Slice Multi-Echo (MSME) sequences with variable TR (TR = 3200 ms, 1600 ms, 800 ms, 400 ms, 200 ms, 100 ms, 50 ms, and 25 ms). T2 maps were acquired using multi-gradient echo (MGE) sequences with variable TE (TE = 10 ms, 20 ms, 30 ms…200 ms). T1 and T2 fitting was performed on the series using ImageJ open-source software (https://imagej.nih.gov/ij/), and a range of T1 and T2 values measured in the tumor were defined. Tumor-applicable T1 values were used to generate a contrast dilution series of gadopentetate dimeglumine (Gd-DTPA; Magnevist®, Bayer HealthCare Pharmaceuticals, Wayne, NJ) in distilled water. T2 values in the tumor were mimicked with a series of agarose concentrations.

Based on observed T1 and T2 values in the tumor, tubes containing a range of Gd-DTPA and agarose concentrations were assembled into a 3D-printed, custom phantom (subsequently referred to as the “study phantom”). A resolution insert was designed for integration into the study phantom which contained laser-cut holes in a 0.5 x 6.0 mm diameter Cirlex disk with holes ranging from 200-75 μm in diameter. The study phantom was loaded with the contrast series and resolution insert, and enclosed in a 20 ml syringe, through which water was pulled and air removed. The phantom was scanned with the surface coil using the described scan protocol (Table II) to assess contrast delineation and spatial resolution. Finally, the bias field of the surface coil array was evaluated by scanning a tube of distilled water and the study phantom. Post-acquisition bias corrections were assessed with the study phantom by placing it in a coil-affixed tube where it was laterally constrained but free to rotate. The coil was positioned and the study phantom was scanned with a series of 2D T1-weighted sequences using the protocol developed for the animal studies. After each scan the phantom was rotated 10-15 degrees. These scans were used to quantitatively measure the efficacy of the bias corrections used in the post-processing pipeline to remove the radiofrequency (rf) sensitivity bias in the 4-element surface coil used for the study.

**Table II.**
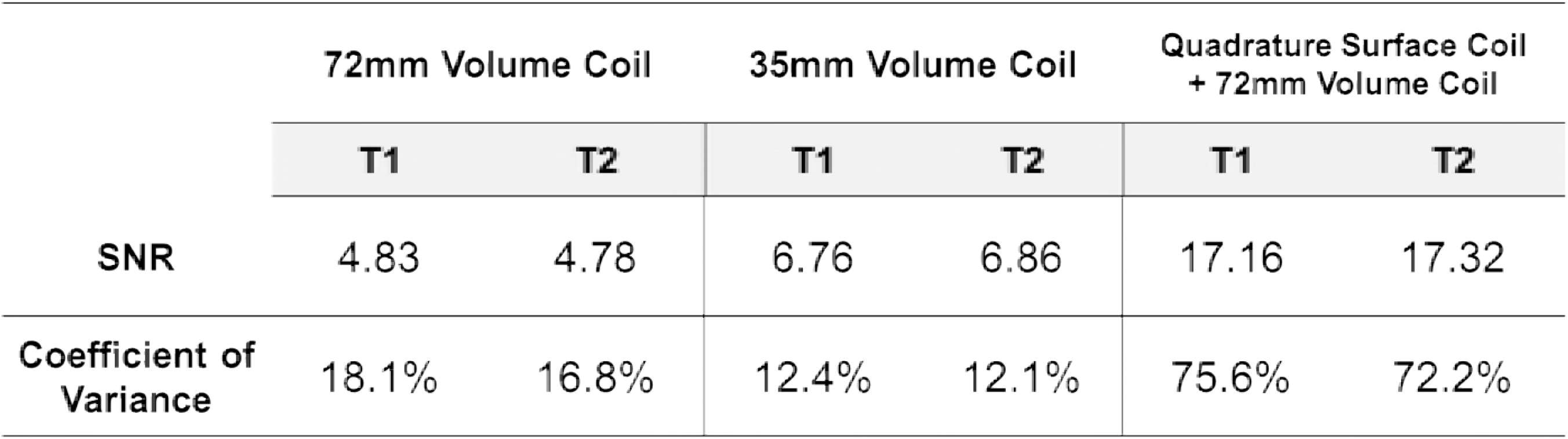
Sensitivity and homogeneity of a uniform sample scanned with three rf coil configurations for small animal MR at 7.0T

#### Mouse MR image acquisition

All animal handling and imaging procedures were performed according to protocols approved by the Duke Institutional Animal Care and Use Committee (IACUC). Tumor images were acquired using the 4-element surface coil array (receive) coupled with the 72 mm linear volume (transmit) coil. Anesthesia was induced with an isoflurane suspension administered on cotton gauze in a contained chamber, followed by maintenance via inhaled isoflurane in concentrations of 1-2% in air. Mice were placed in a left lateral recumbent position on a custom 3D-printed bed equipped with warm water circulation (between 35-40°C) and respiratory rate monitoring. The tumor bearing limb was positioned beneath the surface coil, which was fastened to the coil platform of the bed to reduce displacement. The entire bed platform was positioned within the magnet and mice were monitored for the duration of scanning, with isoflurane concentration adjusted as needed to maintain steady breathing. T1 contrast enhancement was performed by injection of 0.5 mmol/kg Gd-DTPA via tail vein catheter which was placed under anesthesia prior to scanning. Injection speed was 2.5 ml/min, and contrast was allowed to circulate for 3 minutes to achieve peak enhancement. Upon completion, catheters were removed, and mice were returned to warmed cages for anesthesia recovery (<5 minutes).

A systematic comparison of sequences and parameters was undertaken to yield a protocol with T1 and T2 contrast analogous to the clinical trial. Spatial resolution was scaled appropriately for the mouse and tradeoffs between scan parameters balanced to provide signal to noise comparable to the clinical scan in a realistic scan time. The final sequence selections are outlined in Table III. Briefly, the scan protocol was developed which contained a T1-weighted and T2-weighted sequence, followed by repeat of the T1 sequence after Gd-DTPA injection. Upon positioning a mouse in the magnet, routine adjustments were performed, including wobble and shims. Including a 3-minute wait period for circulation of contrast agent post-injection, the entire scan program lasted 40 minutes and 27 seconds for each mouse, plus approximately 5 minutes for initial placement, alignment and adjustments.

**Table III.** Clinically-driven scan program for preclinical MR imaging of soft tissue sarcomas of the extremity

| Sequence | Protocol | Acquisition Parameters |  | Image Geometry |  | Time |
| --- | --- | --- | --- | --- | --- | --- |
| T1-Weighted | T1_FLASH | TE | 4.5 ms | FOV | 28 mm X 28 mm | 09m : 44s |
|  |  | TR | 925.486 ms | Matrix | 280 X 280 |  |
|  |  | Averages | 3 | Resolution | 100 μm |  |
|  |  | Flip angle | 30° | Slice thickness | 300 μm |  |
|  |  |  |  | Slice number | 60 |  |
| T2-Weighted | T2_TurboRARE | TE | 45 ms | FOV | 28 mm X 28 mm | 14m : 59s |
|  |  | TR | 8563.193 ms | Matrix | 280 X 280 |  |
|  |  | Averages | 3 | Resolution | 100 μm |  |
|  |  | RARE | 8 | Slice thickness | 300 μm |  |
|  |  |  |  | Slice number | 60 |  |
| T1-Weighted<br>+ Contrast | T1_FLASH | TE | 4.5 ms | FOV | 28 mm X 28 mm | 03m : 00s<br>+<br>09m : 44s |
|  |  | TR | 925.486 ms | Matrix | 280 X 280 |  |
|  |  | Averages | 3 | Resolution | 100 μm |  |
|  |  | Flip angle | 30° | Slice thickness | 300 μm |  |
|  |  | Start delay | 3 min. post-inj. | Slice number | 60 |  |
|  |  |  |  |  | Total | 37m : 27s |

TE: Echo time TR: Repetition time

#### *In vivo* tumor volume assessment with MR

Volume measurement of tumors imaged with MR was performed with multiple methods to assess accuracy, precision, reproducibility, and inter-user variance. T2-weighted images (as described in Table III) were used in refining and testing volume calculation methods. Slice-by-slice, hand-drawn segmentation was used as the “gold standard” for volume determination, with initial segmentations performed in triplicate to determine user precision. Hand-drawn segmentations were executed in the 2D Viewer platform of OsiriX DICOM viewing software version 9.0.2 (Pixmeo Sarl, Bernex, Switzerland).

Semi-automated tumor segmentation was also explored and refined in 3D Slicer (https://www.slicer.org), an open-source software developed as part of the National Alliance for Medical Image Computing (NA-MIC) under the NIH. Tumors were segmented with a modified protocol for image adjustment and automatic volume propagation. Briefly, DICOM images obtained with the surface coil array were subjected to a bias correction to compensate for signal fall-off which occurs at increasing distances from the coil surface. Bias correction was performed under the N4ITK MRI Bias Correction module in 3D Slicer (45) using the T1-weighted image without contrast as a mask, and the resulting bias-corrected T2 images were used for segmentation. Volume propagation was performed by defining regions of tumor, as well as surrounding non-tumor tissues, on one slice of each orthogonal view near the center of the lesion with the paintbrush editor. Following planar region of interest (ROI) definition, the “GrowCut” tool was used to grow the 3D volume of interest (VOI) considered to be a tumor (46), and the resulting volumes were refined with the “Remove Islands” tool. Semi-automated segmentation was performed in triplicate and results were compared to hand-drawn volumes.

### Micro-CT imaging

All micro-CT imaging was performed using a micro-CT system developed in house (47). Free breathing animals were scanned under anesthesia using 2–3% isoflurane delivered by nose-cone. A pneumatic pillow positioned on the thorax was connected to a pressure transducer to monitor breathing and for respiratory gating. Body temperature was maintained with heat lamps and a feedback controller. Scan parameters were 80 kVp, 40 mA, 10 ms/exposure. A total of 360 views were acquired over a 360° rotation. The reconstruction was performed with a 63 um isotropic voxel size using Feldkamp algorithm (48) followed by bilateral filtration (49) to reduce noise. The micro-CT images were converted to Hounsfield units (HU) and saved as DICOM files. The radiation dose associated with a micro-CT scan was ~0.017 Gy per mouse. This is ~294 to 411 times less than LD50/30 lethal dose (5-7 Gy) in mice (50).

#### Phantom examinations for CT

To assess image quality, a commercially available performance evaluation micro-CT phantom (www.simutec.com) was imaged. The phantom (model vmCT 610) incorporates six plates, each of which is designed to evaluate different aspects of micro-CT image quality with a single scan. These include CT number calibration, CT number linearity, image noise, image uniformity, spatial resolution, geometric accuracy.

#### Micro-CT acquisition using prospective respiratory gating

Unlike in clinical chest CT, which is performed in a single breath hold, preclinical projection data in micro-CT must be acquired over many breaths, requiring respiratory gating. Respiratory gating can be performed prospectively or retrospectively (51). In prospective respiratory gating, a single respiratory phase (e.g. end-expiration) can serve well to assess lung nodules. We were first to develop and implement combined cardiac and respiratory gating for micro-CT (52), which provides the highest possible cardio-pulmonary imaging quality. But for this preclinical study focused on assessing lung metastases longitudinally in a large number of mice, adding cardiac gating substantially increases acquisition times and was considered not essential. Consequently, we have used prospective gating to synchronize acquisition with respiration only using a respiratory signal provided by a pneumatic pillow positioned on the chest of the animal. In prospective gating, the acquisition of each projection is triggered when the respiratory signal crosses above a user-defined threshold. Thus, all projections are acquired in the same part of the respiratory cycle e.g. in end-inspiration, minimizing motion artifacts and blurring in the reconstructions.

To assess the performance of our gated lung tumor imaging, we used mice with primary lung cancer. Lung tumors were generated by intranasal injection of Adeno-Cre into LSL-*Kras^G12D^*; *p53^fl/fl^* mice (53, 54). Mice were imaged at 12 weeks post Adeno-Cre infection, at which point multiple primary lung tumors (~0.5–1.5 mm in diameter) were detectable within each mouse. Each mouse was scanned three times *in vivo* with and without gating. A post-mortem scan was also performed for each mouse and deemed the gold standard for lesion sizes. The lung tumors were semi-automatically segmented using 3D Slicer with the GrowCut tool as previously described for MR images. Volumes were calculated three times per condition, which included a non-gated *in vivo* image, a gated *in vivo* image, and a post-mortem image acquired (standard).

### Development of web-accessible archives for protocol and data dissemination

The Center for In Vivo Microscopy (CIVM) at Duke has initiated a novel approach to sharing data in all of our publications through the use of VoxPort/VoxStation. This integrated package was developed under National Cancer Institute (NCI) support (CA088658) specifically for project management of this nature. VoxPort is a MYSQL database that provides the user with tools for capture and upload of a wide range of data types: IACUC protocols, imaging and set up protocols, 2, 3 and 4D images (more than 30 different formats) from multiple sources (MRI,CT, conventional histology) and data analysis (Excel, graphics, etc). Voxport annotates and organizes these data for efficient search and review by an external user. VoxStation, the companion software, provides external users interactive access to data without requiring download of large, unnecessary files.

### Statistical analysis

Reliability and repeatability of volume measurements were assessed using one-way ANOVA as well as by calculating coefficients of variance, where p<0.05 was considered statistically significant. Differences in output measurements were compared with the student’s t-test, where p<0.05 was considered statistically significant. Precision of repeated measures were interpreted via Brown-Forsythe testing. Statistical analyses were performed and visualized using GraphPad Prism version 7.00 for Mac (GraphPad Software, La Jolla California, US).

## Results

### Establishment of a preclinical protocol for volume measurement of primary sarcomas

Our primary objective was to define a preclinical MR protocol that provided spatial resolution and contrast differential comparable to that of the clinical arm of the trial. To make the protocol practical it must replicate the three acquisitions used in the clinical arm i.e. T1 weighted, T2 weighted, and T1 with contrasts (Fig 1). The final constraint placed on the protocol was that it had to be executed in < 1 hr. Since the mouse is ~ 3000 times smaller than the human, the spatial resolution should be scaled comparably. The spatial resolution in the human protocol (Supporting Table 1) is 1.0 x 2.0 x 5.0 mm i.e. voxels of 10 mm^3^. Thus, our target resolution (voxel volume) is 0.003 mm^3^. The exceptional contrast of MR becomes important in sarcoma lesions receiving RT, as the volume of responding tumors may temporarily increase due to tissue damage and edema, possibly resulting in false assumptions of progression (55, 56). Both T1 and T2 contrast in MR are dependent of the magnetic field as T1 increases and T2 decreases with field (57). Clinical studies are performed at 1.5-3.0 T. This preclinical arm is performed at 7.0 T, so TR and TE have been adjusted iteratively within the rest of the constraints of the protocol to achieve contrast differences between tumor and muscle that are comparable to the clinical arm.

Table II shows the sensitivity and homogeneity for a uniform 18 mm diameter water bottle scanned using the three different rf coil configurations: 72 mm volume coil, transmit receive; 35 mm volume coil, transmit receive; 72 mm volume coil transmit, 4-element surface coil receive. The relative sensitivity of the two volume coils is well defined, with demonstrated homogeneity superior to the surface coil. However, the approximately 3-fold higher sensitivity of the surface coil provides a compelling argument for its use, as desirable SNR is achievable with relatively short scan times. The 5-fold increase in variance of image intensity when using the surface coil is attributable to location-dependent bias. Placement of the surface coil directly onto the tumor-bearing limb reduces the distance between the coil and tumor tissues, somewhat reducing the effect of inhomogeneity in identifying lesions. However, the remaining bias is able to be addressed with the employment of a bias correction (see Fig 4).

3D isotropic imaging is frequently employed in preclinical imaging to provide signal averaging required for the smaller voxels. Clinical MR protocols use 2D (anisotropic) multi-slice sequences to maintain practical scan times. We compared both 2D anisotropic and 3D isotropic sequences in sarcoma-bearing limbs. The contrast to noise ratio and scan times are shown for several variations in (Supporting Fig 1). 2D sequences for T1-weighted and T2-weighted images with 100 um in-plane (axial) resolution and slice thickness of 300 μm were achievable with 3 NEX with short acquisition times (~10 minutes and ~15 minutes, respectively). Further, these images were acquired over a field of view (FOV) sufficient to cover any tumor imaged on study.

In following the direction of the clinical MR program, a T1-weighted and T2-weighted sequence was selected, with T1-acquisition performed without and with injection of contrast. A time course of short T1-weighted acquisitions was performed to identify peak contrast time in tumor tissues (Supporting Fig. 2). The resulting sequences were incorporated into an MR scan protocol suitable for clinically-relevant imaging of primary soft tissue sarcoma lesions of the hind leg for purposes of longitudinal tumor volume assessment (Table III). A comparison of clinical MR images of a patient with a sarcoma in the leg and a tumor-bearing mouse hind limb is shown in Fig 2. Importantly, the clinically-driven preclinical protocol is achievable in less than one hour providing the efficiency necessary for a large study.

**Fig 2.**
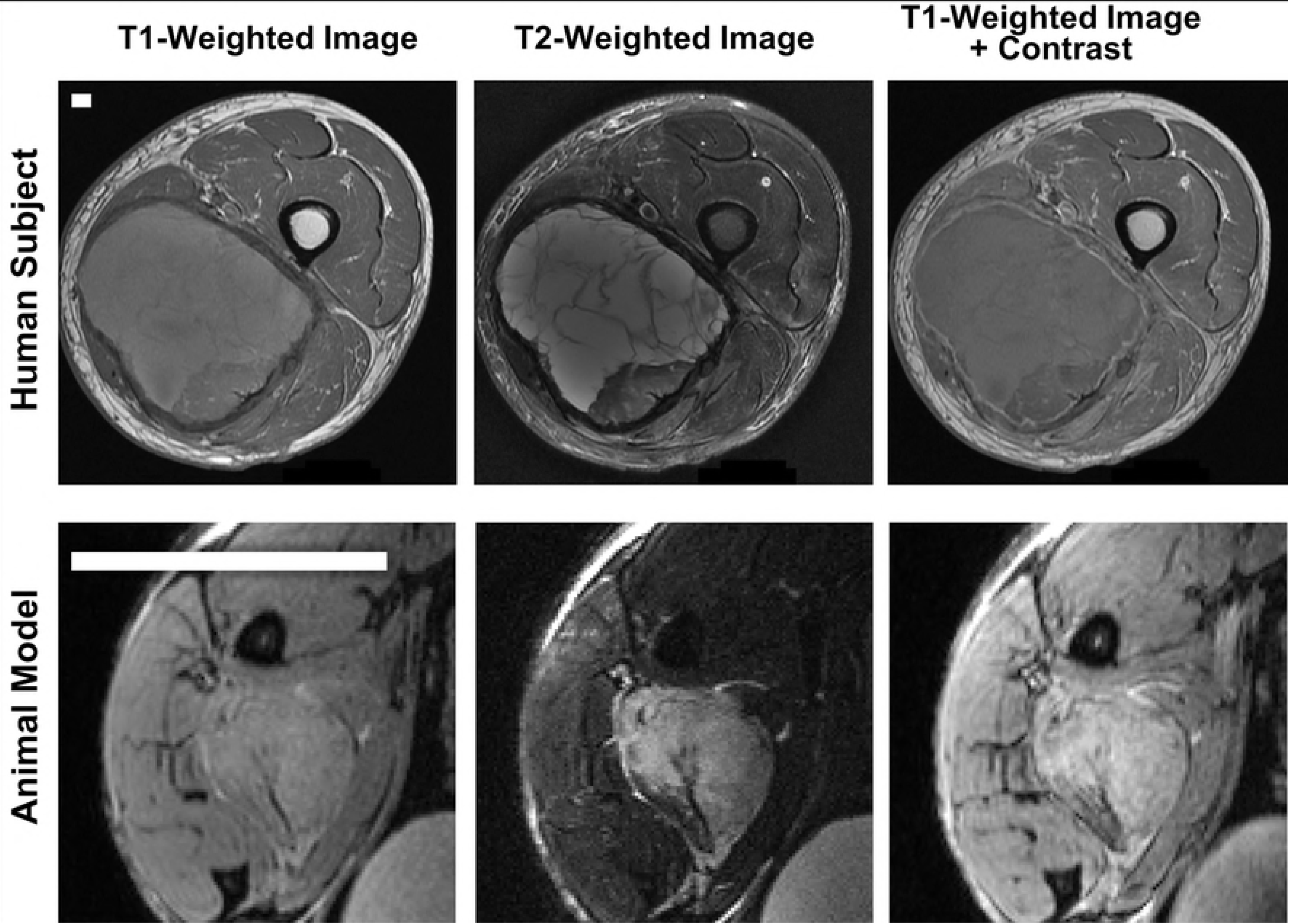
Comparison of human and mouse MR images acquired for the co-clinical trial. Micro-MR images of a sarcoma-bearing mouse leg (bottom row) were obtained with a scan program designed to mimic images acquired in patients enrolled in the clinical arm (top row) bearing soft tissue sarcomas of the extremity. T1-weighted (left), T2-weighted (middle), and T1-weighted + contrast agent injection (right) were acquired in both the human and the mouse arms of the co-clinical trial. White scale bars indicate distances of 5 mm.

### Phantom scans for qualification of preclinical MR systems in achieving a clinically-relevant scan program

Baseline studies were performed as part of a standard quality assurance protocol and validation of scanner performance which served as the preclinical equivalent of standard clinical QA. These protocols were performed regularly throughout study to ensure scanner performance. Scanner linearity, rf coil homogeneity, and magnetic field homogeneity are demonstrated in Supporting Figs 3 and 4, respectively.

Since high-resolution and tissue contrast were the driving motivations for sarcoma imaging with MR, a project-focused “study phantom” was designed to ensure that selected protocols were adequate for tumor volume measurements. To address tissue contrast, T1 and T2 mapping of an established sarcoma were calculated to identify a range of T1 and T2 values which will likely be encountered in tumors on study (Supporting Fig. 5). To mimic contrasts that span the range of tumor-associated T1 values, a series dilution of magnevist was generated. Syringes containing multiple dilutions of magnevist in water were placed in a holder, and T1 mapping was performed to determine the concentration-dependent T1 values. Solutions which produced T1 values within the range present in tumor tissues were selected for incorporation into the study phantom (Fig 3, top). Similarly, a series of agarose concentrations were scanned to determine T2 values, and solutions mimicking T2 values seen in the tumor were selected for phantom construction (Fig 3, bottom). The constructed study phantom was used to confirm the utility of the selected rf coils and scan protocols for successful tumor detection and volume measurement (Fig 4).

**Fig 3.**
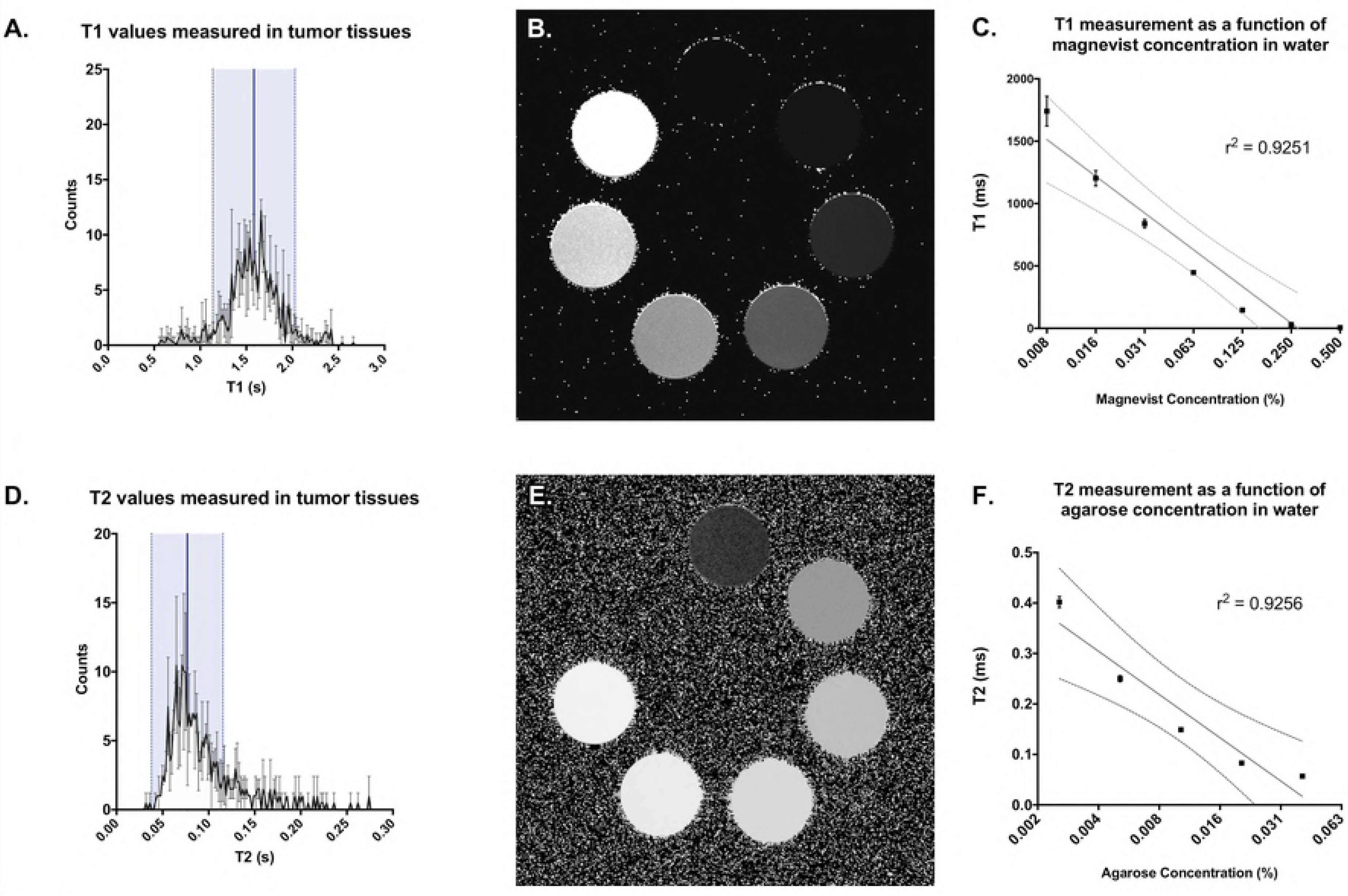
T1 and T2 fitting across a preclinical soft tissue sarcoma demonstrate the anticipated T1 signal range of tumors on study. Histograms of T1 (A), and T2 (D) values measured in sarcoma tissues are shown, including the mean (blue line) and 2 standard deviations (shaded light blue). Bottles containing a dilution series of magnevist (B) and agarose (E) were measured in the 72 mm volume coil and used to mimic the ranges of T1 and T2 values in tumor during construction of a custom study phantom. Linear regressions of T1 or T2 measurements (C and F, respectively) were plotted along a log scale of solution concentration and shown with 95% CI (dotted lines).

**Fig 4.**
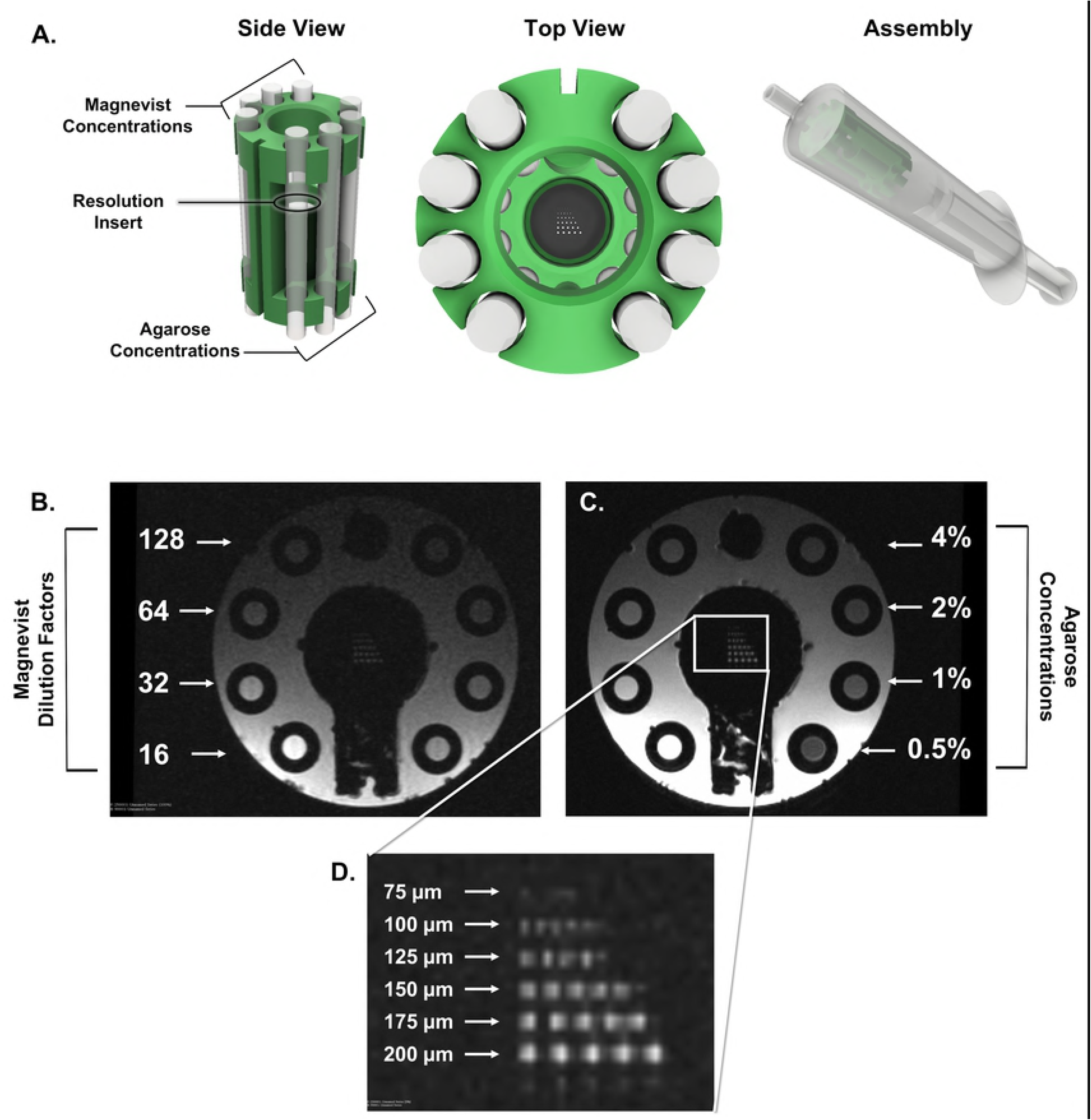
A custom study phantom demonstrates T1 + T2 range and resolution for mouse sarcoma imaging. A 3D-printed phantom was designed to hold tubes containing a range of magnevist (T1) and agarose (T2) concentrations, as well as a resolution insert, and was loaded into a syringe filled with water (A). The T1 sequence used in the preclinical trial demonstrates the range of T1 signal within predetermined magnevist concentrations (B), where dilution factor refers to the dilution of a 1% solution in deionized water. The T2 sequence used in the preclinical trial demonstrates the range of T2 signal within predetermined agarose gel concentrations (C). The resolution insert confirms sufficient resolution of the sequences down to 100um (D). Images shown have not been altered or corrected for bias.

### Evaluation and correction of image biases present during high-field MR imaging

When operating at higher fields, artifacts may appear that are not seen or are negligible in clinical scans. One of the most notable of these inconsistencies is the introduction or amplification of biases in the resulting images. At 7.0 T, the rf wavelength approaches dimensions of the area of interest being imaged, often leading to an area of unexpected brightness in the image center (58-60). This effect, attributed to dielectric resonance, was measured using two uniform phantoms: a 30 ml syringe filled with 10 mM CuSO_4_ and one filled with silicon oil (Supporting Fig. 5). T1-weighted and T2-weighted sequences (defined in Table I) both demonstrated measurable central brightening across the CuSO_4_ phantom, an effect not appearing in the silicon oil (58). Thus, we identified dielectric resonance as a source of bias in the resulting mouse images.

An independent and more obvious bias exists in image intensity as a function of distance from the surface coil itself. The spatial (B1) sensitivity of the multi-coil is a widely recognized bias that is exacerbated at high field strengths. The resulting shifts in signal intensity can confound tumor detection and measurement, particularly in deep-seated tissues. To overcome this issue, we identified the best parameters with which to implement N4ITK bias correction in the study images using 3D Slicer (45, 61). Further improvement of the bias correction was achieved by using a contrast-reduced T1 image (TE = 4.5 ms; TR = 3000 ms), which reflected only the spatial bias, as a weighted filter during correction. Original T1-weighted images of the study phantom demonstrate the overwhelming effect of the surface coil spatial bias, where magnevist concentration dependence is degraded based on distance from the coil surface (Fig 5 A, left). However, application of filtered bias correction (Fig 5 A, right) resulted in restoration of magnevist concentration-dependence in the T1 image signal (Fig 5 B). This served as rationale for application of a bias correction to T2-weighted images on which tumor volumes would be calculated due to better contrast. The scan program includes a T1-weighted image without contrast injection which provides little contrast of use in defining tumor at 7.0 T. However, this can be employed for weighted filtering of the refined N4ITK bias correction, which was subsequently applied to images to improve tumor volume analysis.

**Fig 5.**
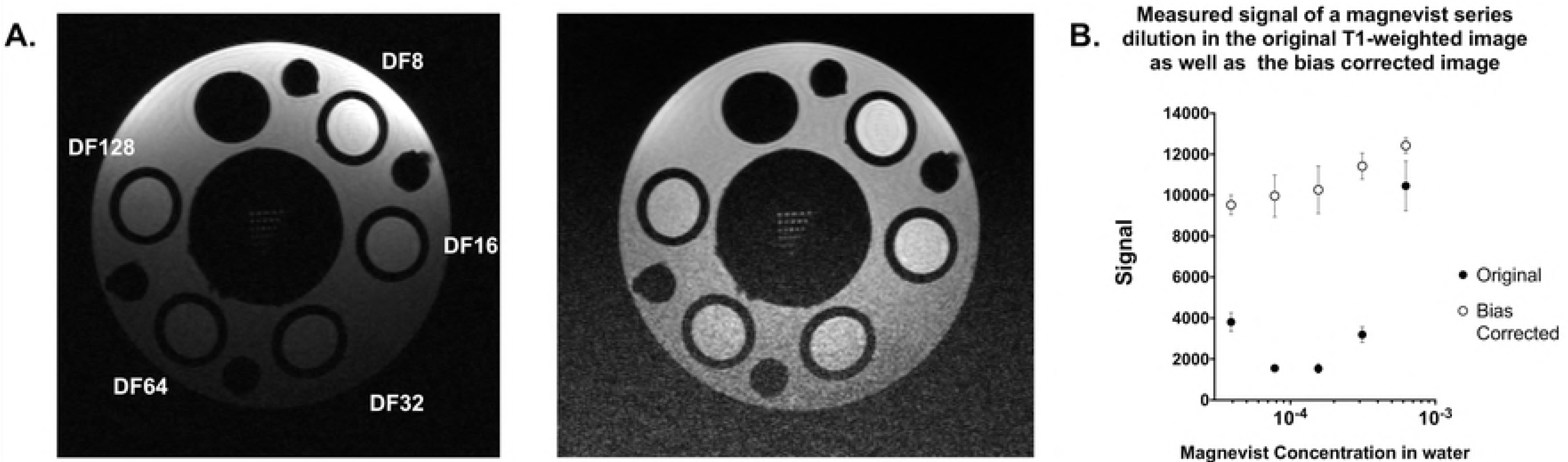
Correction of position-dependent bias resulting from use of the surface coil. A custom phantom containing concentrations of magnevist with T1 values that span those anticipated in tumors were scanned using the surface coil. T1-weighted imaging of the phantom demonstrated a significant bias (A, left), which distorted the intensity of the magnevist tubes due to distance from the coil. Resulting measurements were unable to reflect the concentration-dependent signal appropriately (B). Application of an N4ITK bias correction, masked with a proton density weighted image (TR = 3000) reduced the effects of the coil bias (A, right), restoring linearity in the signal vs concentration curve (B).

### MR scans with the quadrature surface coil deliver reliable and repeatable volume estimates despite repositioning of tumor-bearing limbs

With the goal of reliable tumor volume measurement, images acquired with the clinically-driven MR scan protocol were used to determine its practical utility. First, reliability of scans for volume measurements was tested by scanning tumor-bearing limbs three times in succession with repositioning using the same scan program (Fig 6 A). In each of three scanned mice with morphologically distinct tumors, calculated volumes did not differ based on leg position (Fig 6 C-D). Further, no significant difference in measurement precision was observed in any scan position (Fig 6 E). These data suggest that employment of the clinically-driven MR scan protocol was repeatable and reproducible, regardless of tumor position or orientation beneath the surface coil.

**Fig 6.**
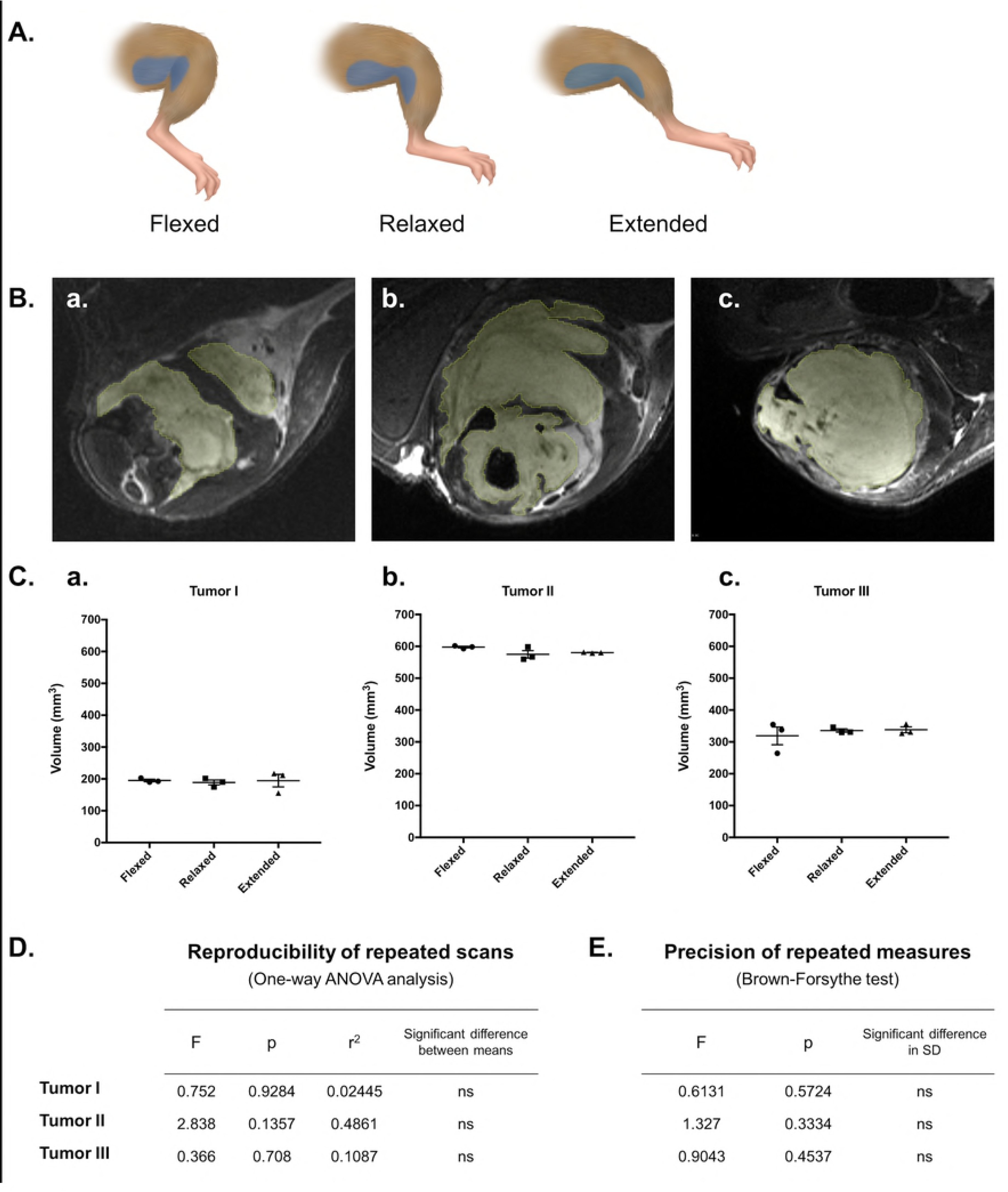
Reliability and consistency of tumor volume measurements in repeated scans with limb repositioning. T2-weighted images were analyzed to determine reproducibility of tumor volume measurements resulting from three consecutive scans of sarcoma-bearing limbs in three positions. Each mouse was scanned three times in succession, with the tumor-bearing limb positioned under the surface coil in a flexed, relaxed, or extended position to shift the location and shape of the tumor (represented as blue in the diagram shown in (A)). ROIs were hand-drawn slice-by-slice in triplicate in each resulting image (9 total measurements per mouse) (B), and calculated volumes were compared for repeatability (user precision) and reproducibility (consistency with shifting position) (C). ANOVA analysis of volume reproducibility suggested no dependence of volume measurements on leg position (D), and precision of hand-drawn measurements was confirmed using Brown-Forsythe (E).

### Semi-automated tumor segmentation demonstrates similar accuracy and precision to hand-drawn measurements of tumor volume, with reasonable inter-user variance

With the inclusion of an advanced bias correction, we have employed tools in 3D Slicer for semi-automated segmentation of T2-weighted MR images to measure tumor volumes efficiently and reliably in a large number of animals. Semi-automated segmentation was compared to volumes calculated with hand-drawn ROIs (“gold-standard”) in T2-weighted tumor images corrected for bias (Fig 7 A and B). ANOVA of the resulting tumor volumes revealed no significant difference in calculated volumes or precision between methods (Fig 7 C). Further, when both techniques were applied to 6 independent tumor image samples, Bland-Altman analysis confirmed reasonable agreement between segmentation methods (Fig 7 D). Taken together these data suggest that volumes calculated by applying a bias correction and subsequent semi-automated tumor segmentation in 3D Slicer are comparable to results from hand-drawn analyses. For studies with large animal numbers, this is very advantageous, as the semi-automated segmentation protocol usually requires <5 minutes to complete.

**Fig 7.**
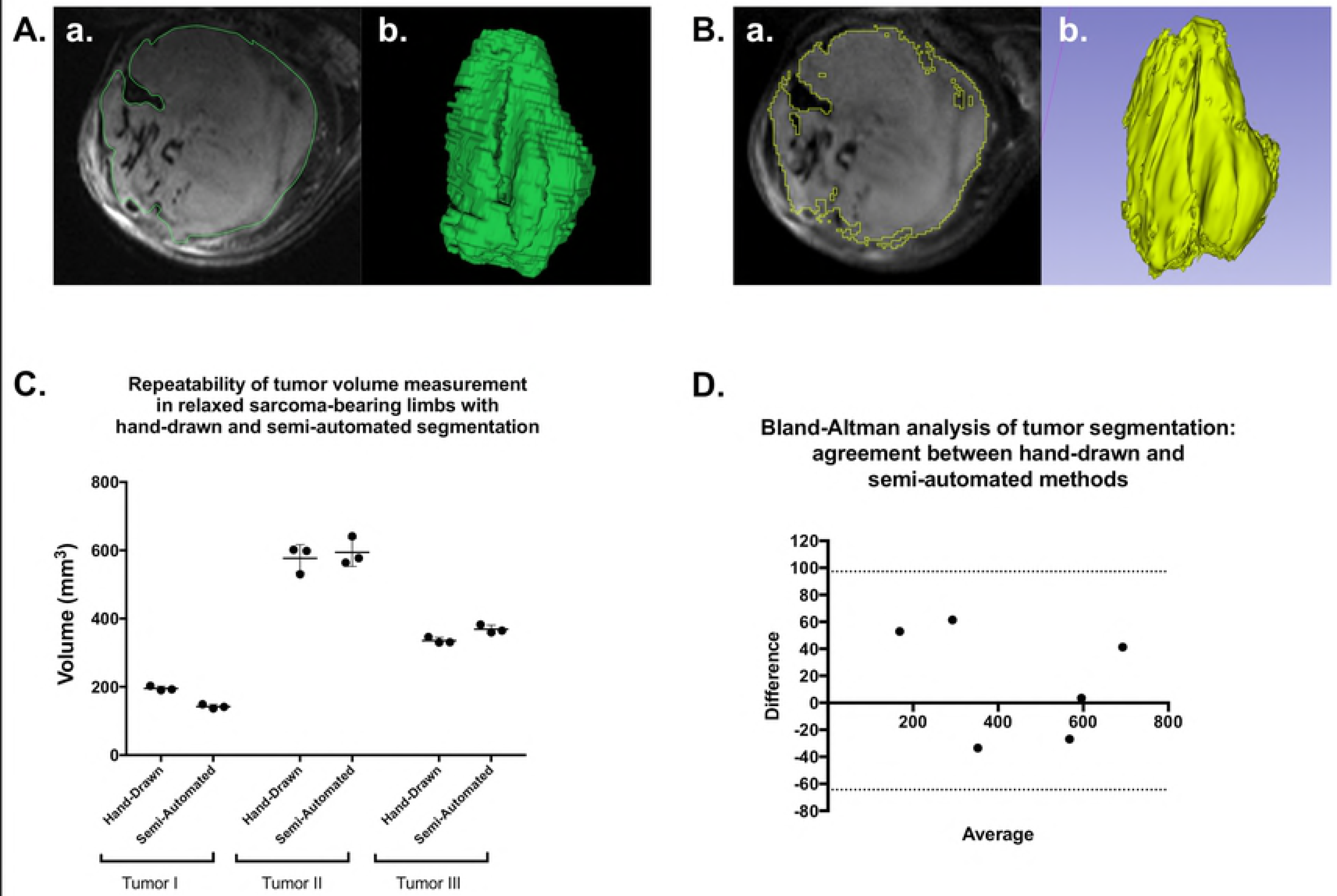
Semi-automated tumor segmentation with 3D Slicer is an acceptable alternative to hand-drawn ROIs for sarcoma volume measurement. T2-weighted images of three tumors of varying size and morphology were analyzed with hand-drawn ROIs and semi-automated segmentation to calculate tumor volume. Examples of hand-drawn (A) and semi-automated (B) segmentation of the same tumor are shown, including an axial slice of the segmented tumor (a.) and the rendered 3D representation of the tumor according to each segmentation (b.). Resulting volume calculations were compared between triplicate measures with each method for each tumor (C), demonstrating similar measurements and comparable precision. Bland-Altman analysis of both methods, when employed in 6 separate tumors, indicated that semi-automated segmentation is a viable alternative to hand-drawn segmentation.

### Employment of respiratory gating during micro-CT acquisition improved tumor burden assessment in lung tissues

Baseline studies were performed as part of a standard quality assurance protocol and validation of scanner performance which served as the preclinical equivalent of standard clinical QA. These protocols were performed regularly throughout study to ensure scanner performance. Spatial resolution, geometric accuracy, iodine concentration measurement, and uniformity were all assessed (Supporting Fig. 6).

During a non-gated micro-CT (Fig 8 A), lung motion often causes blurred tissue boundaries, particularly along the lung wall or near the diaphragm (Fig 8C). Respiratory gating (Fig 8 B) improves the ability to identify small tumors and discern borders between tissues in contact (Fig 8 D).

**Fig 8.**
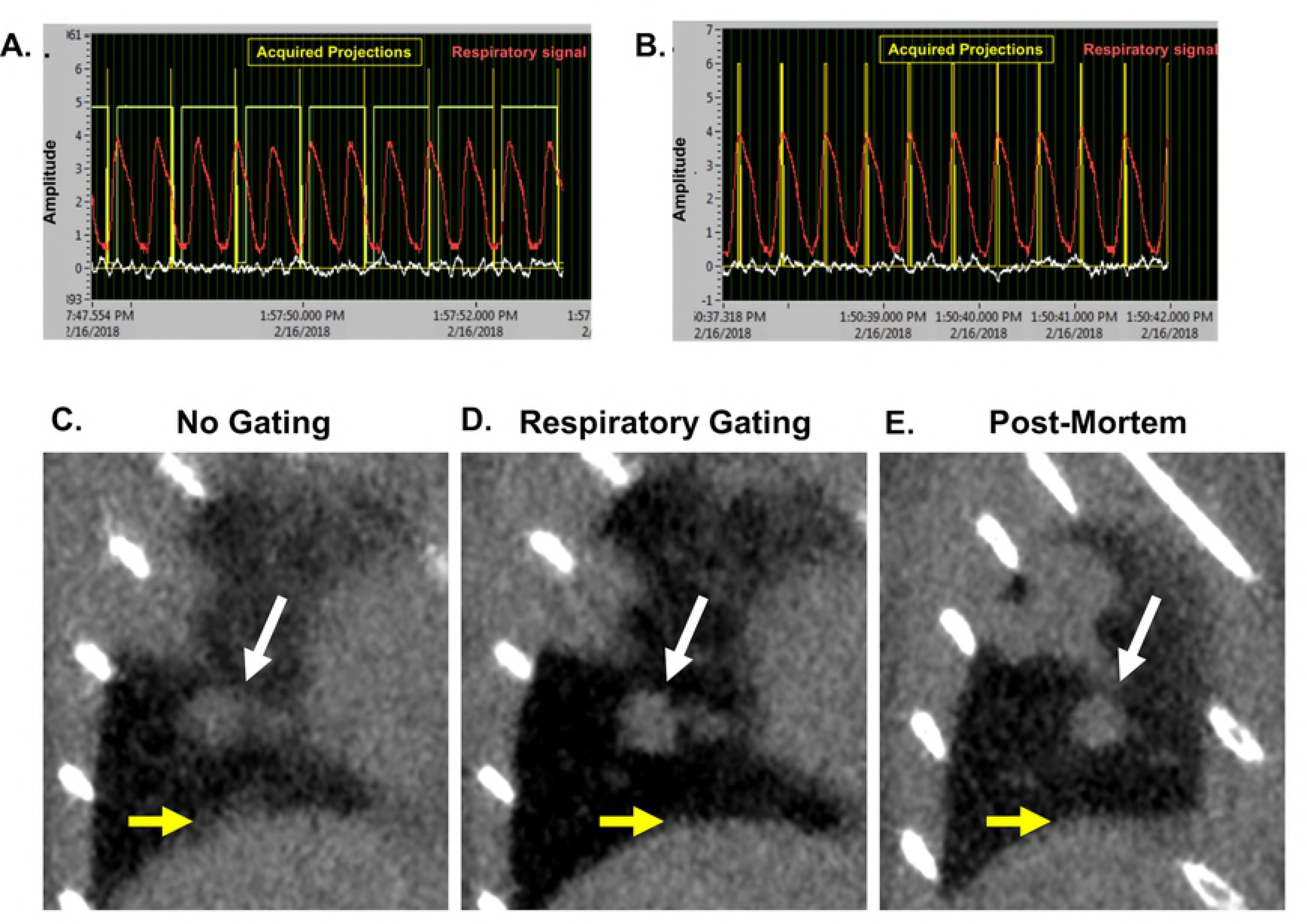
Prospective gating allows for synchronization of image acquisition with breathing patterns. Micro-CT acquisition monitoring without (A) and with respiratory gating (B). Coronal images of a lung tumor are shown without gating (C), with respiratory gating (D), and post-mortem (E). Tumor is indicated by white arrows, and diaphragm is noted with yellow arrows. Representative images from triplicates are shown.

A collection of tumors was selected for identification and volume measurement in each of the 9 acquired images (3 non-gated, 3 respiratory-gated, and 3 post-mortem scans). Selected tumors varied in size and location with diverse surrounding structures (Fig 9A). Mean volume measurements of tumors in respiratory-gated images more closely reflected values measured in post-mortem images than did non-gated tumor measurements. One-way ANOVA of measurements collected for each tumor showed that half of the samples demonstrated significant differences in output values in non-gated images when compared to post-mortem samples (Fig 9B and C). Further, better precision was observed in gated analyses than non-gated analyses (Fig 9D). Taken together, these data support that respiratory gating reduces effects of breathing motion in both the accuracy and precision of lung tumor volume measurements.

**Fig 9.**
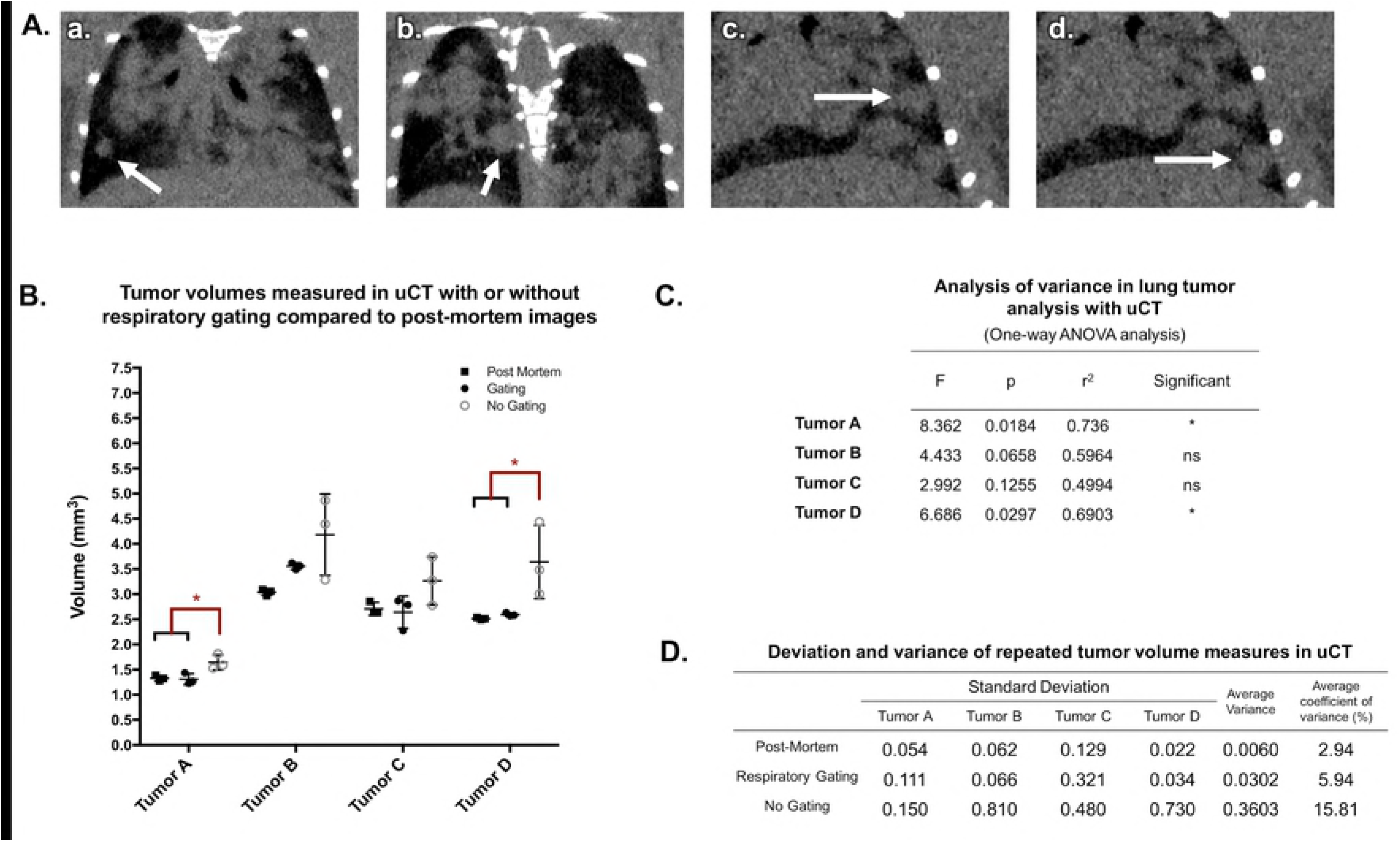
Respiratory gating improves precision and accuracy of lung tumor volume assessment in a free-breathing mouse. Multiple tumors of varying size and location in the lungs of a free-breathing mouse scanned with micro-CT were identified (A, a-d). Hand-drawn ROIs were used to calculate the volume of each lesion in each scan (a total of 9 measurements per tumor). ANOVA showed that in half of the lesions, respiratory gating had a significant impact on the accuracy of volume measurement (B and C). Variance within repeated measures from gated images was approximately double that of post-mortem images, while non-gated image results demonstrated more than five-fold variance compared to post-mortem measurements (D).

### Databases constructed via VoxPort allow users web-access to co-clinical animal imaging data, acquisition and analysis protocols, and supporting materials

This integrated package was developed under NCI support (CA088658) specifically for project management of this nature. VoxPort is a MYSQL database that provides the user tools for capture and upload of a wide range of data types: IACUC, imaging and set up protocols, 2,3 and 4D images (more than 30 different formats), from multiple sources (MRI,CT, conventional histology) and data analysis (Excel, graphics, etc). Voxport annotates and organizes these data so an external user can efficiently search and review. VoxStation, the companion software, provides external users interactive access to all these data without need to download vast troves of what might be useless.

Fig 10 illustrates Voxport/Voxstation in action displaying both images and protocols. Access to our Voxport Image Management System is available at: https://civmvoxport.vm.duke.edu/voxbase/index.php. The visitor will be prompted to login to CIVMVoxPort using the provided credentials (User Name: DukeU24, Password: DukeU24Review)”.

**Fig 10.**
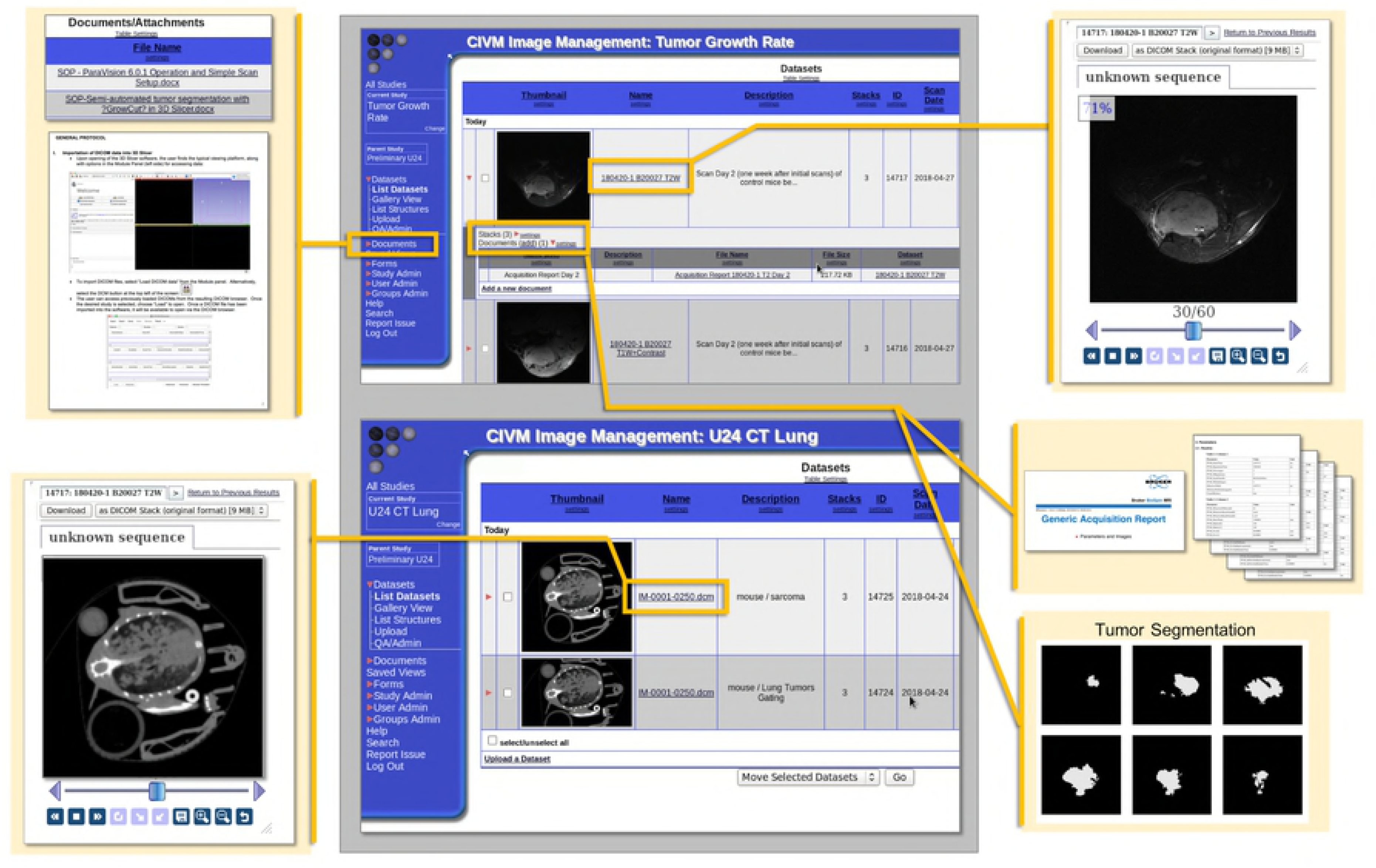
Demonstration of the VoxPort interface for image data sharing. Images of VoxPort demonstrating archives of multiple forms of imaging data, including images from multiple modalities, protocol documentation and standard operating procedures, scan acquisition information, and segmentation stacks which correspond to archived images. The image gallery provides a thumbnail and metadata about the images. The user can choose to download a given image or stack or examine the data within the interface.

## Discussion

In order to bridge the gap between preclinical and clinical research, care must be taken to develop animal models and imaging protocols that appropriately reflect the clinical question. To serve a co-clinical study of PD-1 inhibition for sarcoma, we have described the establishment, qualification, and application of preclinical MRI and CT imaging protocols for longitudinal assessment of therapeutic efficacy in a GEMM of soft tissue sarcoma. The imaging protocols are clinically-driven, reproducible, suitable for high-throughput studies, and readily extended to experiments using different tumor models or interventions.

In selecting micro-MRI protocols, we gained higher spatial resolution (100 μm in-plane) by using a quadrature surface coil placed over the tumor-bearing limbs of animals. We assembled a scan program to generate images similar to those acquired in the clinic (T1-weighed, T2-weighed, and T1-weighted plus contrast) with spatial resolution scaled for the size of the mouse, contrast comparable to the clinical arm of the study, and total acquisition time lasting less than 1 hour. With this program, tumor volume measurements were reliable regardless of tumor size or depth, and independent of shifts in the tumor orientation/position beneath the coil.

Similar to the clinic, periodic imaging of mouse lung tissues following treatment and resection of primary sarcomas is being used to monitor metastatic development. The principal challenge in assessing tumor burden in the lungs with micro-CT is the effect of respiratory motion on resulting images. We have employed respiratory gating techniques to mitigate the effects of motion by limiting acquisition to defined periods in the breathing cycle. Tissue barriers, such as diaphragm, lung wall, and tumors boundaries, were visibly more clear in gated images compared to images acquired throughout the full breathing cycle. As a result, tumor volume measurements from gated images more accurately matched post-mortem standards and demonstrated improved precision compared to measurements from non-gated images, improving tumor burden assessment.

Finally, we have created a web accessible repository to store and share these protocols, representative data, supporting studies (e.g. pathology, blood chemistry). In sharing our findings over the course of the study, we hope to assist others in generating and evaluating their preclinical and co-clinical imaging studies. Overall, we have devised a strategy for incorporation of preclinical imaging into a co-clinical trial for the evaluation of tumor burden, as well as established a means by which to efficiently organize and share the resulting data.

## Conclusions

We have established a routine pipeline by which high-volume preclinical imaging data can support and inform in real time a co-clinical trial of combined RT and immunotherapy in sarcoma. Additionally, we have outlined a blueprint for navigating and overcoming some of the technical challenges that translational tumor imaging studies face. The resulting methods are both clinically-relevant and widely adaptable. Although the field of animal imaging lacks standards of practice and reporting, we have generated a means by which our data, protocols, and processing methods can be accessed and used as a template by others. By establishing clinically-driven preclinical imaging methods to serve in a co-clinical trial, we have created a pipeline which reduces the gap between preclinical and clinical studies of sarcoma therapy.

## Acknowledgements

All small animal imaging work was performed at the Duke Center for In Vivo Microscopy. We thank Lixia Liu and Nerissa Williams with assistance with caring for mice.

## Supporting information

**S1 Table. Comparison of imaging parameters between the clinical and preclinical arms of the co-clinical trial.**

**S1 Figure. 2D tumor imaging achieves adequate signal with reduced acquisition time compared to 3D acquisition.** T2-weighted images of a primary sarcoma lesion of the hind limb were acquired with both 2D (A) and 3D (B) TurboRARE sequences. TE was fixed at 45 ms, and both resulting images had an in-plane resolution of 100um (C). Signal in both images was adequate for deciphering tumor location and boundaries at 1 NEX, but the acquisition time to generate the 3D image was more than 7X that of the 2D acquisition. Reduced scan time of the 2D sequence allows time for incorporation of multiple NEX while maintaining a clinically-relevant scan program length.

**S2 Figure. Magnevist contrast enhancement peaks at approximately 6 minutes post-injection.** A series of short (~ 1 min) T1 FLASH sequences in the hind limb of a tumor-bearing mouse were performed before, during, and after injection of magnevist. Time course images (A) were obtained to determine wash-in, peak, and wash out of contrast in tumor compared to baseline (B). The highest intensity of contrast in tumor was observed approximately 6 minutes following injection. Subsequent scan programs included a delay between injection and scan initiation to align peak contrast within the acquisition window of the T1 protocol.

**S3 Figure. Scanner linearity and rf coil homogeneity.** A large, custom-printed 3D grid phantom comprised of 1 mm rods in 10mM CuSO_4_ was used to demonstrate scanner linearity over the area in which the surface coil is employed (A). Surface coil operation occurs within the defined yellow box. To measure homogeneity, a uniform bottle containing distilled water was scanned with the 72 mm volume coil (B) and the surface coil operating in the 72 mm coil (C), and signal was plotted as a function of distance over the volume of the bottle.

**S4 Figure. Identification of image bias due to dielectric effects measured in images acquired with the 72 mm volume coil.** To measure the influence of the dielectric effect in images acquired using the 72 mm volume coil, images of 30 ml bottles containing 10mM CuSO_4_ in deionized water (A.a.) and Silicone Oil (A.b.) were acquired with the volume coil (transceiver; independent of the surface coil). A linear region of interest with a width of 20 pixels (yellow box) was used to measure signal intensity across the diameter of the bottle. Signal intensity shifts due to the dielectric effect are measurable across the diameter of the CuSO_4_ water bottle (B.a.), and are not present in images of Silicone Oil (B.b.).

**S5 Figure. T1 and T2 maps of sarcoma tumors provide anticipated T1 and T2 values of tumor tissues.** T1 and T2 maps were acquired over multiple slices of an established sarcoma of the hind limb. A T2-weighted TurboRARE sequence was used to delineate tumor ROIs (A). Tumor tissue and anatomical landmarks are indicated (T = tumor; Bl = bladder; S = spine). The tumor ROI used to measure contrast is shown on the T1 map (B) and T2 map (C).

**S6 Figure. Optimization and calibration of the microCT.** Demonstration of the selected phantom (A) as well as image quality evaluation results of micro-CT scans are shown. An axial micro-CT slice thorough the spatial resolution plate with the 4 coils of alternating aluminum and Mylar sheets demonstrate measured layer thicknesses of 50, 100, 150, and 200 μm corresponding to 10, 5, 3.3, and 2.5 line pairs per mm (lp/mm), respectively (B). Only the first 2 coils show spatial frequencies that can be separated, indicating image resolution around 3.3 lp/mm. The phantom contains a geometric accuracy plate which includes two peripheral beads and a central bead. The two measured distances are reasonably similar to the ones indicated by the manufacturer, with an error around 50 μm. This error is under one pixel size value of 63μm (C). A plot of the measured CT number versus known iodine concentration (D.a.; yellow rings) demonstrates a significant linear correlation between signal intensity and increasing iodine concentration (D.b.). A line profile (D.a.; red line) confirms uniformity with relative variations less than 11 % across the field of view without significant cupping artifacts due to beam hardening (D.c.).

## References

1. Ledford H. Translational research: 4 ways to fix the clinical trial. Nature. 2011;477(7366):526-8.

2. Ruggeri BA, Camp F, Miknyoczki S. Animal models of disease: pre-clinical animal models of cancer and their applications and utility in drug discovery. Biochem Pharmacol. 2014;87(1):150-61.

3. Wartha K, Herting F, Hasmann M. Fit-for purpose use of mouse models to improve predictivity of cancer therapeutics evaluation. Pharmacol Ther. 2014;142(3):351-61.

4. Clohessy JG, Pandolfi PP. Mouse hospital and co-clinical trial project--from bench to bedside. Nat Rev Clin Oncol. 2015;12(8):491-8.

5. Lunardi A, Pandolfi PP. A co-clinical platform to accelerate cancer treatment optimization. Trends Mol Med. 2015;21(1):1-5.

6. Chen Z, Cheng K, Walton Z, Wang Y, Ebi H, Shimamura T, et al. A murine lung cancer co-clinical trial identifies genetic modifiers of therapeutic response. Nature. 2012;483(7391):613-7.

7. Nardella C, Lunardi A, Patnaik A, Cantley LC, Pandolfi PP. The APL paradigm and the "co-clinical trial" project. Cancer Discov. 2011;1(2):108-16.

8. Nishino M, Sacher AG, Gandhi L, Chen Z, Akbay E, Fedorov A, et al. Co-clinical quantitative tumor volume imaging in ALK-rearranged NSCLC treated with crizotinib. Eur J Radiol. 2017;88:15-20.

9. Haris M, Yadav SK, Rizwan A, Singh A, Wang E, Hariharan H, et al. Molecular magnetic resonance imaging in cancer. J Transl Med. 2015;13:313.

10. Kauppinen RA, Peet AC. Using magnetic resonance imaging and spectroscopy in cancer diagnostics and monitoring: preclinical and clinical approaches. Cancer Biol Ther. 2011;12(8):665-79.

11. Lin G, Keshari KR, Park JM. Cancer Metabolism and Tumor Heterogeneity: Imaging Perspectives Using MR Imaging and Spectroscopy. Contrast Media Mol Imaging. 2017;2017:6053879.

12. Clark DP, Lee CL, Kirsch DG, Badea CT. Spectrotemporal CT data acquisition and reconstruction at low dose. Med Phys. 2015;42(11):6317-36.

13. Li XF, Zanzonico P, Ling CC, O’Donoghue J. Visualization of experimental lung and bone metastases in live nude mice by X-ray micro-computed tomography. Technol Cancer Res Treat. 2006;5(2):147-55.

14. Day CP, Merlino G, Van Dyke T. Preclinical mouse cancer models: a maze of opportunities and challenges. Cell. 2015;163(1):39-53.

15. Le Magnen C, Dutta A, Abate-Shen C. Optimizing mouse models for precision cancer prevention. Nat Rev Cancer. 2016;16(3):187-96.

16. O’Connor JPB. Cancer heterogeneity and imaging. Semin Cell Dev Biol. 2017;64:48-57.

17. Bokacheva L, Ackerstaff E, LeKaye HC, Zakian K, Koutcher JA. High-field small animal magnetic resonance oncology studies. Phys Med Biol. 2014;59(2):R65-R127.

18. Gillies RJ, Bhujwalla ZM, Evelhoch J, Garwood M, Neeman M, Robinson SP, et al. Applications of magnetic resonance in model systems: tumor biology and physiology. Neoplasia. 2000;2(1-2):139-51.

19. Mowery YM, Ballman KV, Riedel RF, Brigman BE, Attia S, Meyer CF, et al. SU2C-SARC032: A phase II randomized controlled trial of neoadjuvant pembrolizumab with radiotherapy and adjuvant pembrolizumab for high-risk soft tissue sarcoma. Journal of Clinical Oncology. 2018;36(15_suppl):TPS11588-TPS.

20. Fite BZ, Wong A, Liu Y, Mahakian LM, Tam SM, Aina O, et al. Magnetic resonance imaging assessment of effective ablated volume following high intensity focused ultrasound. PLoS One. 2015;10(3):e0120037.

21. Vormoor B, Knizia HK, Batey MA, Almeida GS, Wilson I, Dildey P, et al. Development of a preclinical orthotopic xenograft model of ewing sarcoma and other human malignant bone disease using advanced in vivo imaging. PLoS One. 2014;9(1):e85128.

22. Sapi J, Kovacs L, Drexler DA, Kocsis P, Gajari D, Sapi Z. Tumor Volume Estimation and Quasi-Continuous Administration for Most Effective Bevacizumab Therapy. PLoS One. 2015;10(11):e0142190.

23. Jimenez-Gonzalez M, Plaza-Garcia S, Arizeta J, Bianchessi S, Trigueros C, Reese T. A longitudinal MRI study on lymph nodes histiocytosis of a xenograft cancer model. PLoS One. 2017;12(7):e0181043.

24. Linxweiler J, Korbel C, Muller A, Jungel E, Blaheta R, Heinzelmann J, et al. Experimental imaging in orthotopic renal cell carcinoma xenograft models: comparative evaluation of high-resolution 3D ultrasonography, in-vivo micro-CT and 9.4T MRI. Sci Rep. 2017;7(1):14249.

25. Hectors SJ, Jacobs I, Strijkers GJ, Nicolay K. Multiparametric MRI analysis for the identification of high intensity focused ultrasound-treated tumor tissue. PLoS One. 2014;9(6):e99936.

26. Jeon TY, Kim CK, Kim JH, Im GH, Park BK, Lee JH. Assessment of early therapeutic response to sorafenib in renal cell carcinoma xenografts by dynamic contrast-enhanced and diffusion-weighted MR imaging. Br J Radiol. 2015;88(1053):20150163.

27. Kersemans V, Cornelissen B, Allen PD, Beech JS, Smart SC. Subcutaneous tumor volume measurement in the awake, manually restrained mouse using MRI. J Magn Reson Imaging. 2013;37(6):1499-504.

28. Larkin TJ, Canuto HC, Kettunen MI, Booth TC, Hu DE, Krishnan AS, et al. Analysis of image heterogeneity using 2D Minkowski functionals detects tumor responses to treatment. Magn Reson Med. 2014;71(1):402-10.

29. Ramasawmy R, Johnson SP, Roberts TA, Stuckey DJ, David AL, Pedley RB, et al. Monitoring the Growth of an Orthotopic Tumour Xenograft Model: Multi-Modal Imaging Assessment with Benchtop MRI (1T), High-Field MRI (9.4T), Ultrasound and Bioluminescence. PLoS One. 2016;11(5):e0156162.

30. Whisenant JG, Sorace AG, McIntyre JO, Kang H, Sanchez V, Loveless ME, et al. Evaluating treatment response using DW-MRI and DCE-MRI in trastuzumab responsive and resistant HER2-overexpressing human breast cancer xenografts. Transl Oncol. 2014;7(6):768-79.

31. Zhang F, Cao J, Chen X, Yang K, Zhu L, Fu G, et al. Noninvasive Dynamic Imaging of Tumor Early Response to Nanoparticle-mediated Photothermal Therapy. Theranostics. 2015;5(12):1444-55.

32. Zhang P, Wang B, Chen X, Cvetkovic D, Chen L, Lang J, et al. Local Tumor Control and Normal Tissue Toxicity of Pulsed Low-Dose Rate Radiotherapy for Recurrent Lung Cancer: An In Vivo Animal Study. Dose Response. 2015;13(2):1559325815588507.

33. Almeida GS, Panek R, Hallsworth A, Webber H, Papaevangelou E, Boult JK, et al. Pre-clinical imaging of transgenic mouse models of neuroblastoma using a dedicated 3-element solenoid coil on a clinical 3T platform. Br J Cancer. 2017;117(6):791-800.

34. Bagci U, Kramer-Marek G, Mollura DJ. Automated computer quantification of breast cancer in small-animal models using PET-guided MR image co-segmentation. EJNMMI Res. 2013;3(1):49.

35. Liebsch L, Kailayangiri S, Beck L, Altvater B, Koch R, Dierkes C, et al. Ewing sarcoma dissemination and response to T-cell therapy in mice assessed by whole-body magnetic resonance imaging. Br J Cancer. 2013;109(3):658-66.

36. Rajendran R, Huang W, Tang AM, Liang JM, Choo S, Reese T, et al. Early detection of antiangiogenic treatment responses in a mouse xenograft tumor model using quantitative perfusion MRI. Cancer Med. 2014;3(1):47-60.

37. Yang SH, Lin J, Lu F, Han ZH, Fu CX, Lv P, et al. Evaluation of antiangiogenic and antiproliferative effects of sorafenib by sequential histology and intravoxel incoherent motion diffusion-weighted imaging in an orthotopic hepatocellular carcinoma xenograft model. J Magn Reson Imaging. 2017;45(1):270-80.

38. Blasiak B, Landry J, Tyson R, Sharp J, Iqbal U, Abulrob A, et al. Molecular susceptibility weighted imaging of the glioma rim in a mouse model. J Neurosci Methods. 2014;226:132-8.

39. Dassler K, Scholle FD, Schutz G. Dynamic gadobutrol-enhanced MRI predicts early response to antivascular but not to antiproliferation therapy in a mouse xenograft model. Magn Reson Med. 2014;71(5):1826-33.

40. Muller A, Jagoda P, Fries P, Graber S, Bals R, Buecker A, et al. Three-dimensional ultrashort echo time MRI and Short T2 images generated from subtraction for determination of tumor burden in lung cancer: Preclinical investigation in transgenic mice. Magn Reson Med. 2018;79(2):1052-60.

41. Beloueche-Babari M, Jamin Y, Arunan V, Walker-Samuel S, Revill M, Smith PD, et al. Acute tumour response to the MEK1/2 inhibitor selumetinib (AZD6244, ARRY-142886) evaluated by non-invasive diffusion-weighted MRI. Br J Cancer. 2013;109(6):1562-9.

42. Mustafi D, Fernandez S, Markiewicz E, Fan X, Zamora M, Mueller J, et al. MRI reveals increased tumorigenesis following high fat feeding in a mouse model of triple-negative breast cancer. NMR Biomed. 2017;30(10).

43. Song Y, Cho G, Suh JY, Lee CK, Kim YR, Kim YJ, et al. Dynamic contrast-enhanced MRI for monitoring antiangiogenic treatment: determination of accurate and reliable perfusion parameters in a longitudinal study of a mouse xenograft model. Korean J Radiol. 2013;14(4):589-96.

44. Zhang T, Zhang F, Meng Y, Wang H, Le T, Wei B, et al. Diffusion-weighted MRI monitoring of pancreatic cancer response to radiofrequency heat-enhanced intratumor chemotherapy. NMR Biomed. 2013;26(12):1762-7.

45. Tustison NJ, Avants BB, Cook PA, Zheng Y, Egan A, Yushkevich PA, et al. N4ITK: improved N3 bias correction. IEEE Trans Med Imaging. 2010;29(6):1310-20.

46. Vezhnevets V, Konouchine V. Growcut: Interactive multi-label N–D image segmentation by cellular automata. Proc GraphiCon. 2005;7.

47. Badea C, Johnston S, Johnson B, De Lin M, Hedlund LW, Johnson GA. A Dual Micro-CT System for Small Animal Imaging SPIE, Medical Imaging; San Diego, CA: SPIE; 2008.

48. Feldkamp LA, Davis LC, Kress JW. Practical cone-beam algorithm. J Opt Soc Am. 1984;1(6):612-19.

49. Tomasi C, Manduchi R. Bilateral filtering for gray and color images. Proc of the 1998 IEEE International Conference on Computer Vision. 1998:839-46.

50. Carlson SK, Classic KL, Bender CE, Russell SJ. Small animal absorbed radiation dose from serial micro-computed tomography imaging. Mol Imaging Biol. 2007;9(2):78-82.

51. Badea C, Drangova M, Holdsworth D, Johnson G. In vivo small-animal imaging using micro-CT and digital subtraction angiography. Phys Med Biol. 2008;53::R319.

52. Badea CT, Hedlund LW, Johnson GA. Micro-CT with respiratory and cardiac gating. Medical Physics. 2004;31(12):3324-9.

53. Johnson L, Mercer K, Greenbaum D, Bronson RT, Crowley D, Tuveson DA, et al. Somatic activation of the K-ras oncogene causes early onset lung cancer in mice. Nature. 2001;410(6832):1111-6.

54. Kirsch DG, Grimm J, Guimaraes AR, Wojtkiewicz GR, Perez BA, Santiago PM, et al. Imaging primary lung cancers in mice to study radiation biology. Int J Radiat Oncol Biol Phys. 2010;76(4):973-7.

55. Cuneo KC, Mito JK, Javid MP, Ferrer JM, Kim Y, Lee WD, et al. Imaging primary mouse sarcomas after radiation therapy using cathepsin-activatable fluorescent imaging agents. Int J Radiat Oncol Biol Phys. 2013;86(1):136-42.

56. Messiou C, Bonvalot S, Gronchi A, Vanel D, Meyer M, Robinson P, et al. Evaluation of response after pre-operative radiotherapy in soft tissue sarcomas; the European Organisation for Research and Treatment of Cancer-Soft Tissue and Bone Sarcoma Group (EORTC-STBSG) and Imaging Group recommendations for radiological examination and reporting with an emphasis on magnetic resonance imaging. Eur J Cancer. 2016;56:37-44.

57. Johnson GA, Herfkens RJ, Brown MA. Tissue relaxation time: in vivo field dependence. Radiology. 1985;156(3):805-10.

58. Bernstein MA, Huston J, 3rd, Ward HA. Imaging artifacts at 3.0T. J Magn Reson Imaging. 2006;24(4):735-46.

59. Collins CM, Liu W, Schreiber W, Yang QX, Smith MB. Central brightening due to constructive interference with, without, and despite dielectric resonance. J Magn Reson Imaging. 2005;21(2):192-6.

60. Hoult DI, Phil D. Sensitivity and power deposition in a high-field imaging experiment. J Magn Reson Imaging. 2000;12(1):46-67.

61. Fedorov A, Tuncali K, Fennessy FM, Tokuda J, Hata N, Wells WM, et al. Image registration for targeted MRI-guided transperineal prostate biopsy. J Magn Reson Imaging. 2012;36(4):987-92.

